# MOVIE: Multi-Omics Visualization of Estimated contributions

**DOI:** 10.1101/379115

**Authors:** Sean D. McCabe, Dan-Yu Lin, Michael I. Love

## Abstract

**Summary:** The growth of multi-omics datasets has given rise to many methods for identifying sources of common variation across data types. The unsupervised nature of these methods makes it difficult to evaluate their performance. We present MOVIE, Multi-Omics Visualization of Estimated contributions, as a framework for evaluating the degree of overfitting and the stability of unsupervised multi-omics methods. MOVIE plots the *contributions* of one data type against another to produce *contribution plots*, where contributions are calculated for each subject and each data type from the results of each multi-omics method. The usefulness of MOVIE is demonstrated by applying existing multi-omics methods to permuted null data and breast cancer data from The Cancer Genome Atlas. Contribution plots indicated that principal components-based Canonical Correlation Analysis overfit null data, while Sparse multiple Canonical Correlation Analysis and Multi-Omics Factor Analysis provided stable results with high specificity for both the real and permuted null datasets.

**Availability:** MOVIE is available as an R package at https://github.com/mccabes292/movie

**Supplementary information:** Supplementary data are available at *Bioinformatics* online.

## 1 Introduction

With the availability of multi-omics datasets, an increasing number of methods have been developed to decipher the relationship among large scale data types. Some methods, such as iCluster+ (Shen et al, 2010) and Similarity Network Fusion (SNF) (Wang et al, 2014), classify samples into groups, such as tumor subtypes. Other methods determine which features or biological processes contribute to the common variation across all data types, as well as the magnitude of their contributions. These methods include Sparse multiple Canonical Correlation Analysis (Sparse mCCA) (Witten and Tibshirani, 2009), Angle-based Joint and Individual Variation Explained (AJIVE) (Feng et al, 2018), and Multi-Omics Factor Analysis (MOFA) (Argelaguet et al, 2017). Additionally, Canonical Correlation Analysis (CCA) (Hotelling, 1936) can be modified for a high-dimensional setting by running the analysis on the top principal components (PCs) of each matrix. Due to the unsupervised nature of these methods, assessment of performance, in terms of the stability of the output and the degree of overfitting, can be difficult. We propose MOVIE as a tool to evaluate method performance by-examining the contribution of each sample in each data type towards the common variation space and utilizing a k-fold cross validation to assess stability and potential overfitting.

## 2 Methods

Each of the four aforementioned methods decomposes sources of common variation across data types. CCA identifies weights for each feature to maximize the correlation between two separate weighted matrices. In analyses where the datasets have a large number of features, CCA can be conducted on the top PCs of each matrix. Unfortunately, CCA is not generalizable to settings with more than two matrices, and a consensus has not been reached on the appropriate number of PCs to be used. In null data, correlations as high as 0.9 are possible when the number of PCs included in the analysis is large (Supplementary Fig. 1). Sparse mCCA addresses these issues by imposing a sparsity parameter to identify only the relevant features and incorporate multiple large-dimensional matrices. MOFA and AJIVE are two methods that decompose the variation as common across datasets or unique to the individual dataset. AJIVE utilizes matrix perturbation theory to decompose the variation structure across and within data types, while MOFA is a Bayesian factor analysis method that identifies latent factors explained by some or all of the data types.

All of the above methods provide sets of weights corresponding to the importance of each feature in each data type. The larger the absolute value of the weights, the more the feature contributes to the common variation. Except for Sparse mCCA, each method provides multiple sets of weights (whereas Sparse mCCA provides only-one set by default). MOVIE only considers the top set of weights to evaluate method performance (Supplementary Fig. 2). By multiplying the weights by the observed values of each data type, we can obtain *contributions* for each sample and data type combination (Supplementary Methods 1.1 and Supplementary Fig. 3). Plotting contributions of one data type against another allows the user to visualize the strength of the relationships between data types. Additionally, samples that fall off the diagonal in a contribution plot may be biologically meaningful outliers. We call this plot the *contribution plot* and use this tool to evaluate these methods via data splitting. Data splitting and the projection of estimated contributions was proposed by Soneson et al (2010) for parameter tuning and validation of a multi-omics method. Other methods have used leave-one-out cross-validation and the projection of learned factors on new datasets to assess method performance (Brown et al, 2018; Fertig et al, 2012). By-omitting a subset of the samples from the analysis and predicting their contributions from each data type in the training set, we can discern whether the relationships from the full analysis suffer from overfitting or provide unstable results. Using the results of multi-omics methods, scaled contributions can be calculated to make the contribution plot.

The first step in utilizing MOVIE is to conduct a k-fold cross-validation analysis and a full analysis using the selected method. Our software package provides template code in the vignette for performing this step for each method. The number of folds must be chosen such that the number of samples per fold is at least 20, and careful consideration must be taken when selecting fold membership to ensure balanced folds (Supplementary Methods 1.2). The cross-validation analysis is then performed by analyzing the samples in the training set and calculating contributions for those in the test set.

Because the results of many methods may differ slightly in scale across fold, while still identifying the same components of variation, MOVIE scales the contributions within each fold to compare contributions across folds on the same scale (See Supplementary Methods 1.3). Scaling helps with small variations in estimation, but it will not address potential issues related to the identification of different components of the variation space in separate folds. After scaling, contribution plots can be constructed for both the cross-validated and full analyses, and comparisons can be made between the two plots. Additionally, MOVIE provides *comparison plots* that plot the contributions from the cross-validated analysis to the contributions from the full analysis for a specific data type. This plot can be used to evaluate whether the results of these methods are stable with the exclusion of a subset of the samples.

## 3 Results

To demonstrate the usefulness of MOVIE, we constructed contribution and comparison plots for Sparse iriCCA, A JIVE, and MOFA on both real and simulated datasets. We applied these three methods to 558 breast, cancer samples from The Cancer Genome Atlas (TCGA) (Wong et al, 2017). Copy number variation was measured for 216 genes: RNA expression was measured for 12,434 genes: and miRNA expression was measured for 305 genes. Five folds were selected for the analysis, and fold membership was determined by stratifying the samples with the first PC of RNA expression. Supplementary Figs. 4-6 provide side-by-side contribution plots for Sparse iriCCA constructed by MOVIE and indicate that the method did not overfit. The comparison plot shows that the sample contributions maintained their ordering in the cross-validated and full analyses, indicating the consistency of Sparse iriCCA with the removal of samples from the analysis (Supplementary Fig. 7). Plots indicated consistency for MOFA (Supplementary Figs. 8-11) and less consistency for AJIVE (Supplementary Figs. 12-15), although the latter could be due to the identification of different components of the variation space in separate folds.

Using the same TCGA dataset as before, we randomized sample order for both the RNA and miRNA datasets to create a null dataset. The above three methods, along with PC-CCA, were analyzed to determine which methods returned a false-positive relation. Only RNA and miRNA expression data were analyzed to allow for the inclusion of PC-CCA. Fig. 1a shows the side-by-side contribution plots for Sparse iriCCA and demonstrates correctly that there is no relation between the two data types in the null set. Fig. 1b shows a strong linear relation in the full analysis and no relation in the cross-validated analysis for PC-CCA, thus demonstrating that PC-CCA greatly overfits the data and that MOVIE can identify such overfitting. MOFA and AJIVE correctly identified a null result, (Supplementary Figs. 16-19).

**Figure 1:**
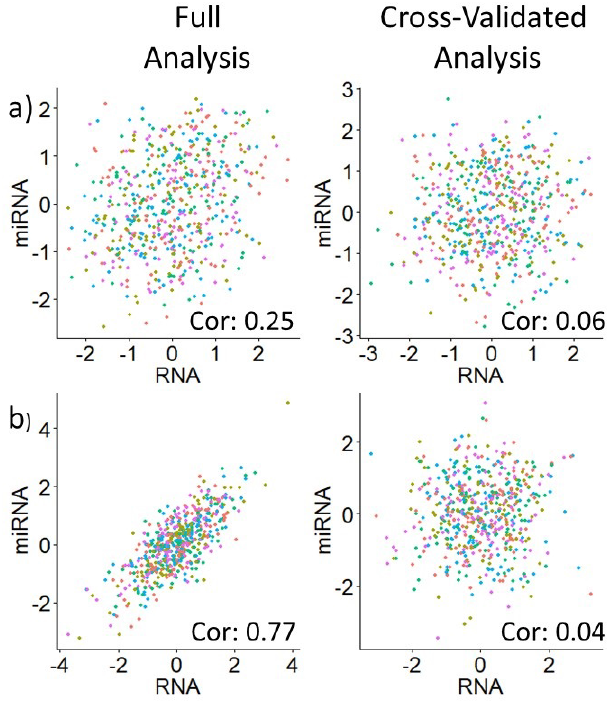
Side-By-Side Contribution Plots for a) Sparse mCCA in null data and b) PC-CCA in null data: Left, panels are the contribution plots from the full analysis, while right, panels are the contribution plots for the cross-validated analysis. Points are colored by the fold membership of each sample.

Due to the unsupervised nature of multi-omics methods for variance decomposition, determining a preferred method is difficult,. Using a data,-split,ting approach, MOVIE provides a framework to compare the performance of unsupervised multiomics methods through the construction of the contribution plot.

